# Dynamics between horizontal gene transfer and acquired antibiotic resistance in *S*. Heidelberg following *in vitro* incubation in broiler ceca

**DOI:** 10.1101/684787

**Authors:** Adelumola Oladeinde, Kimberly Cook, Steven M. Lakin, Zaid Abdo, Torey Looft, Kyler Herrington, Gregory Zock, Jodie Plumblee Lawrence, Jesse C. Thomas, Megan S. Beaudry, Travis Glenn

## Abstract

The chicken gastrointestinal tract harbors taxa of microorganisms that play a role in the health and disease status of the host. The cecum is the part of the gut that carries the highest microbial densities, has the longest residence time of digesta and is a vital site for urea recycling and water regulation. Therefore, the cecum provides a rich environment for bacteria to horizontally transfer genes between one another via mobile genetic elements such as plasmids and bacteriophages. In this study, we used broiler chicken cecum as a model to investigate antibiotic resistance genes that can be transferred *in vitro* from ceca flora to *Salmonella enterica* serovar Heidelberg (*S*. Heidelberg). We used whole genome sequencing and resistome enrichment to decipher the interactions between *S*. Heidelberg, gut microbiome and acquired antibiotic resistance. After 48 h incubation of ceca under microaerophilic conditions, one *S*. Heidelberg isolate was recovered with an acquired Inck2 plasmid (88 kb) encoding extended β-lactamase producing gene (*bla*_CMY-2_). *In vitro*, this plasmid was transferrable between *E. coli* and *S*. Heidelberg strains, but transfer was unsuccessful between *S*. Heidelberg strains. An in-depth genetic characterization of transferred plasmids suggests that they share significant homology with P1-like phages. This study contributes to our understanding of the dynamics between an important food-borne pathogen and the chicken gut microbiome.

**Importance:** *S.* Heidelberg is a clinically important serovar, linked to food-borne illness and among the top 5 serovars isolated from poultry in USA and Canada. Acquisition of new genetic material from microbial flora in the gastrointestinal tract of food animals, including broilers, may contribute to increased fitness of pathogens like *S.* Heidelberg and may increase their level of antibiotic tolerance. Therefore, it is critical to gain a better understanding on the dynamic interactions that occur between important pathogens and the commensals present in the animal gut and other agroecosystems. In this study, we show that the native flora in the broiler ceca were capable of transferring mobile genetic elements carrying AmpC β-lactamase (*bla*_CMY-2_) gene to an important food-borne pathogen *S*. Heidelberg. The potential role for P1-like bacteriophage transduction was also discussed.

## Introduction

Numerous studies have demonstrated that horizontal gene transfer (HGT) plays an important role in the spread of antimicrobial resistance (AMR) and virulence genes to food-borne bacterial pathogens (1). The notion that these genes are horizontally transferred from commensal bacteria residing in the animal gut or environment to food-borne pathogens is becoming more recognized. Likewise, plasmid-mediated antibiotic resistance gene (ARG) transfer is thought to occur between intestinal bacteria and putative pathogens like *Salmonella*. In an earlier study, we reported that *S*. Heidelberg strains that acquired Col-like plasmids from poultry litter microbiota survived longer and demonstrated an increased resistance to selected antibiotics (2). Gumpert et al. also demonstrated that *Escherichia coli* (*E. coli*) strains present in the gut of infants could acquire multidrug (MDR) resistance plasmids in the absence of antibiotic treatment (3). *Salmonella* Heidelberg is one of the top *Salmonella* serovars causing food-borne illness in USA and Canada (4, 5) and poultry has been reported as the primary source (5, 6).

*Enterococcus faecalis* on the other hand is part of the normal microbiome of poultry and present in high abundance in day-old chicks (7). However, *E. faecalis* can be an opportunistic pathogen causing diseases in poultry and humans (8–10). Only a few studies (11, 12) have investigated their promiscuity to plasmid-mediated ARG of poultry origin, consequently our knowledge on their fitness factors in the chicken gut is limiting.

In this study, we investigated the dynamics between HGT and acquired antibiotic resistance in *S*. Heidelberg following *in vitro* inoculation into broiler ceca. Our primary aim was to determine ARG transfer events that occurred after inoculation of *S*. Heidelberg into the chicken ceca, rather than mimicking what would have happened *in vivo*. We show that AmpC β-lactamase gene (*bla*_CMY-2_) carried on an IncK2 plasmid could be transferred successfully between *E. coli* and *S*. Heidelberg but transfer between *S*. Heidelberg strains was not possible. Our analysis also supports the notion that phage-mediated transfer of MDR plasmids is ubiquitous and should be considered in models predicting AMR transfer rates across bacterial lineages.

## Materials and methods

### Bacterial isolates

A strain of *Salmonella enterica* serovar Heidelberg (SH-2813) isolated from chicken carcass rinsate was spontaneously made resistant to 200 ppm of nalidixic acid (nal) (SH-2813_nal_) (2). This resulted in two isogenic *S*. Heidelberg strains with different *gyrA* substitutions conferring nal resistance. One strain had a Ser83tyr substitution while the other was a Ser83phe substitution. These strains are referred to as SH-2813-Parental^tyr^ and SH-2813-Parental^phe^, respectively. The parental SH-2813 strain carries a ∼37 kb IncX1 conjugative plasmid (2). *Enterococcus faecalis* strain JH2-2 (*E. faecalis*_JH2-2_) with chromosomally conferred resistance to rifampicin, fusidic acid and lincosamides was obtained from Dr. Charlene Jackson (USDA-ARS, Athens, GA). The JH2-2 strain harbors no plasmid.

### Bacterial inoculum preparation

Frozen bacterial stocks of SH-2813_nal_ (SH-2813-Parental^tyr^ and SH-2813-Parental^phe^) and *E. faecalis*_JH2-2_ were recovered on XLT-4 and enterococcosel (BD Difco, Sparks, MD) agar supplemented with 200 ppm and 512 ppm of nalidixic acid (nal) and rifampicin (rif) (Sigma-Aldrich Corp, St. Louis, MO), respectively. Plates were incubated at 37 °C for 48 h and 6 colonies were randomly selected for inoculation into Brain Heart Infusion Broth (BHIB) with no antibiotics. After inoculation, BHIB tubes were incubated in a water bath overnight at 37 °C and 85 rpm. Following overnight growth, BHIB culture was diluted 20-fold with fresh BHIB and incubated for ∼3 h in a water bath shaker at 37 °C and 200 rpm. Afterwards, bacterial cells were centrifuged at 4,600 X *g* for 5 mins and washed in 1X Phosphate Buffer Saline (PBS). Washed cells were used as inoculum.

### Ceca inoculation

Entire viscera were collected randomly from 18 broiler birds from the evisceration line of a commercial processing plant. Viscera were put in individual whirl-pak bags and transported immediately on ice for processing. Ceca were removed from viscera and the open ends of one cecum/bird was injected with ∼10^5^ colony forming units (CFU) of SH-2813_nal_ and *E*. *faecalis*_JH2-2_ (Fig. 1). Following inoculation, cecum was immersed in an equal weight volume of sterile 1X PBS and incubated at 37 °C under microaerophilic conditions (5 % O_2,_ 10 % CO_2_, 85 % N_2_) for 48 h.

**Fig. 1.**
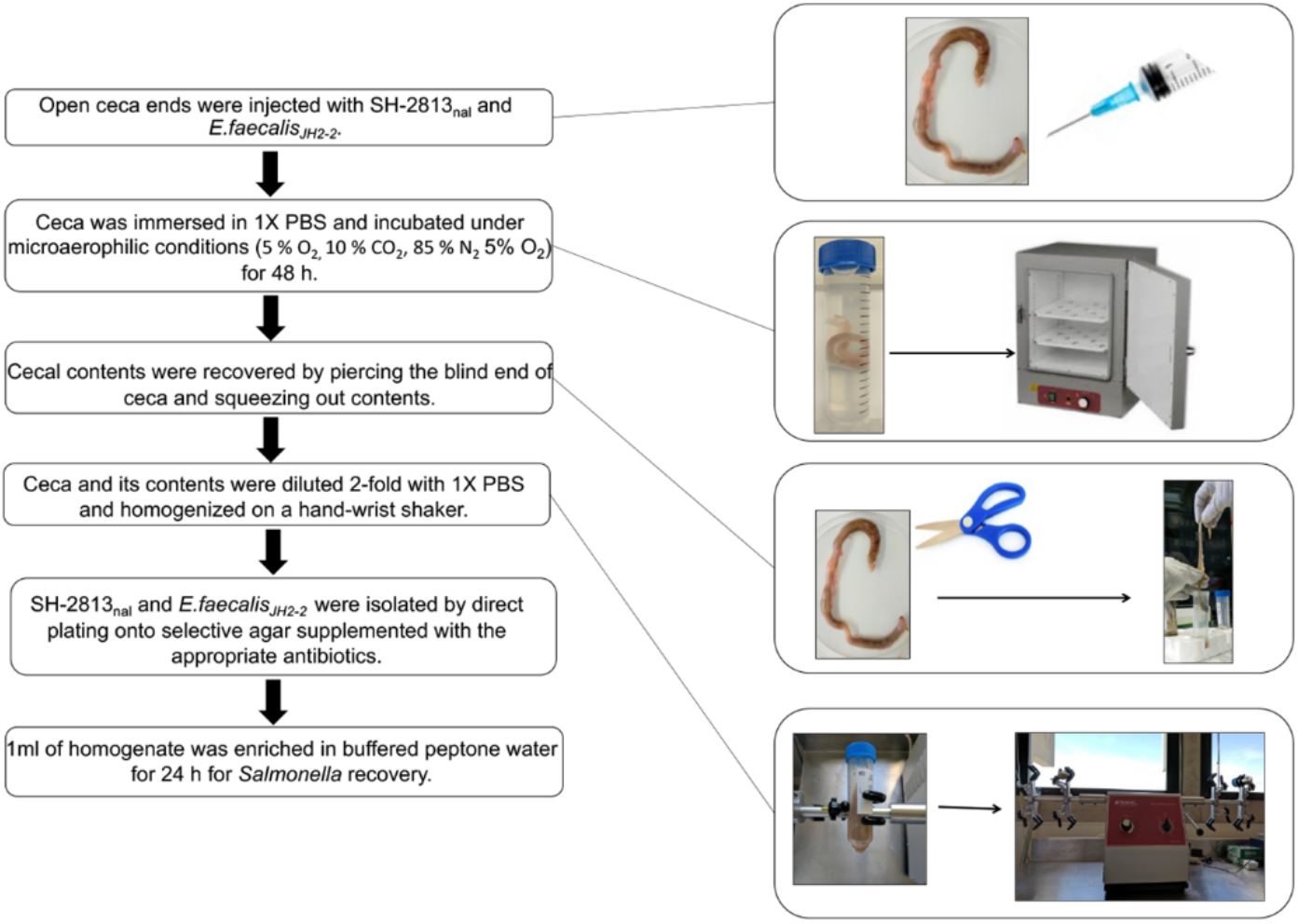
Experimental design. Entire viscera were collected from 18 broiler birds from the evisceration line of a commercial processing plant in Athens, GA, USA. Viscera were put in individual whirl-pak bags and transported immediately on ice to the USDA-ARS research unit in Athens, GA, USA. Ceca were removed from viscera and the open ends of one cecum/bird was injected with SH-2813_nal_ and *E*. *faecalis*_JH2-2_. Ceca was incubated at 37 °C under microaerophilic conditions for 48 h.

### Microbiological analysis

After ceca inoculation, three individual cecum were sampled independently at 0.5, 6, 24 and 48 h. Cecal contents were removed by piercing the blind end of the cecum with sterile scissors and squeezing out contents with sterile gloves. Afterwards, ceca and its contents were diluted 2-fold with 1X PBS and homogenized in a hand-wrist shaker at 450 rpm for 5 mins. Nalidixic resistant *S*. Heidelberg were recovered by direct plating of serial dilutions on XLT-4 agar supplemented with 200 ppm of nal (XLT-4_nal_). Serial dilutions were also performed on XLT-4 agar supplemented with 200 ppm nal and 32 ppm ampicillin (amp) (XLT-4_nal+amp_) for isolation of *S*. Heidelberg clones with acquired resistance to amp. For *E. faecalis*_JH2-2_, rifampicin (rif) resistant isolates were recovered on Enterococcosel agar supplemented with 512 ppm of rif (Enterococcosel_rif_). For selection of JH2-2 clones with acquired resistance to erythromycin (ery), serial dilutions were screened on enterococcosel agar supplemented with 512 ppm of rif and 8 ppm of ery (Enterococcosel_rif+ery_). Ampicillin and erythromycin were chosen to represent the common classes of broad-spectrum β-lactam and macrolide drug resistance found in *Salmonella* serovars and enterococci strains. We also quantified the total population of *E. coli* and Enterococci in uninoculated ceca at pre-inoculation (n = 3) and at 48 h (n = 3) using CHROMagar^TM^ ECC and enterococcosel agar with no antibiotics. All bacterial incubation was carried out at 37°C, unless otherwise noted. The contents of the ceca were saved at a 1:1 ratio in RNAlater (Thermofisher, Waltham, MA) at −80 °C before DNA was extracted or further analysis.

### Antibiotic susceptibility testing

Antimicrobial susceptibility testing was performed on selected SH-2813_nal_ (n = 13) and *E. faecalis*_JH2-2_ (n = 9) isolates recovered during the 48 h incubation of ceca and on isolates from *in vitro* mating experiments (n= 14) using the National Antimicrobial Resistance Monitoring System (NARMS) protocol (13). In addition, a Gram-Negative panel (GN2F) (Thermo Fisher Scientific, Waltham, MA) was used to determine the minimum inhibitory concentrations (MIC) of one isolate to β-lactam drugs that are not available on the NARMS panel. MIC for isolates were determined by broth microdilution using the Sensititre semi-automated antimicrobial susceptibility system (Thermo Fisher Scientific, Waltham, MA). Results were interpreted according to Clinical and Laboratory Standards Institute (CLSI) guidelines when available (CLSI M100-S27); otherwise, breakpoints established by NARMS were used (13).

### Whole genome sequencing and single nucleotide polymorphism identification

Sequencing was performed on SH-2813_nal_ (n =27) isolates recovered from cecal contents after incubation of ceca and on selected SH-2813_rec_ (n = 6) and EC_rec_ (n = 1) strains that acquired antibiotic resistance during *in vitro* mating experiments. WGS was performed as described previously (2). A single run of sequencing on the MiSeq platform resulted in low read depth coverage for some samples, so a second set of technical replicates were sequenced using the same methodology for each sample.

Single nucleotide polymorphisms (SNPs) and indels present in sequenced isolates were determined by aligning raw sequence reads to the chromosome of *S*. Heidelberg strain AMR588-04-00435 (NC_CP016573) using Burrows-Wheeler Aligner (BWA) (14). To avoid false positive SNPs from sequencing or batch effect error, only the sequence reads from the second technical replicates were used in this analysis. SAM file sorting and removal of PCR duplicates was done using SAMtools (15). Genome Analysis ToolKit (16) v 3.6 with a minimum mapping quality of 30 and a minimum base quality of 30 was used for SNP identification. Manipulation of generated variant call format files (VCF) was done using vcftools v 0.1.12b (17).

### Resistome analysis

Cecal DNA was used for resistome analysis. DNA was extracted and purified from 250 mg of cecal +RNAlater homogenate using a previously described method for poultry litter (2). This DNA was used for target enrichment of shotgun libraries to characterize the antibiotic resistance genes (i.e. the resistome) present in ceca. Shotgun metagenomic libraries were prepared using New England BioLabs Ultra FS II library kits (New England BioLabs, Ipswich, MA), ligated to iTru adapter stubs (18), cleaned with a 0.8 ratio of speedbeads after ligation, and amplified with iTru dual-indexing primers. Following library preparation, samples were cleaned with a 1:1 ratio of speedbeads, quantified with a Qubit 2.0 Fluorometer DNA high sensitivity assay kit (Thermofisher, Waltham, MA), and pooled in sets of 8 samples for target enrichment by hybridization capture. Baits were designed and synthesized by Arbor Bioscience (Ann Arbor, MI) for all ARG’s present in the Comprehensive Antibiotic Resistance Database (CARD) (24) database on November 2016. Sequencing was done on the Illumina MiSeq platform with 250-bp paired end reads using the MiSeq reagent V2 (500 cycles).

### Bioinformatics

Whole genome sequence reads from technical replicates were combined and assembled de novo into contigs using SPAdes and plasmidSPAdes (19, 20). The quality of assembled genomes of combined runs was assessed using Quality Assessment Tool for Genome Assemblies (QUAST) (21) (File S1). Assembled contigs were submitted to the Center for Genomic Epidemiology’s ResFinder (22) and CARD for the identification of resistance genes encoded on plasmids or chromosome (23). Contigs were also submitted to PlasmidFinder (24) to determine existing plasmid replicon types.

Prophage-like regions were identified using PHAST (25) and PHASTER (26), and putative sequences were extracted for annotation. Annotation was done with Prokka (27) and by performing a PSI-BLAST search against the NCBI non-redundant (nr) database for viruses. Phylogenetic tree reconstruction of plasmids was performed using the maximum likelihood method implemented in RAxML-NG v. 0.6.0 (28) with the number of bootstrap replicates criteria set to 100. We used the best model of sequence evolution predicted by jModelTest for tree reconstruction (29). ProgressiveMAUVE (30) and MAFFT (31) implemented in Geneious Prime v. 2019.1.1 was used for the comparative analysis of plasmids and phages.

Fastq reads for resistome libraries were assembled using SPAdes (`--meta`). Contigs with ≤10X coverage or shorter than ∼500 bp were not used for ARG identification. Contigs were queried against ResFinder for acquired ARG determination and against CARD for global transcriptional regulators H-NS. The relative abundance of an ARG (log2 fold-change) was determined from the coverage of the contig carrying the ARG, normalized against the coverage of H-NS or Crp. Prior to using this method on the resistome dataset, we benchmarked it against WGS data of susceptible and MDR strains of *S*. Heidelberg (n= 1), *S*. Kentucky (n= 4), *S*. Enteritidis (n= 1), *Campylobacter jejuni* (n= 3), *Campylobacter coli* (n= 4) and *E. faecalis* (n= 4) (see supplemental methods).

### Identification of *E. coli* strains carrying IncK2 plasmids

One hundred microliter of cecal+RNAlater homogenate from time point 48 h (n =6) was spread on to CHROMagar and incubated overnight. Afterwards, 16 *E. coli* colonies were selected randomly and re-streaked on fresh CHROMagar plates. Hot boil DNA extraction (2) was done on pure colonies before screening with two qPCR primers targeting IncK2 (Table S1).

**Table 1.** Bacterial population in inoculated broiler ceca ^a^.

| Time (h) | Bacterial concentration (Log CFU/g of cecum)<br>Mean (SD) |  |  |  |  |
| --- | --- | --- | --- | --- | --- |
|  | SH-2813<sub>nal</sub> <sup>c</sup> | *E. faecalis*<sub>JH2-2</sub> <sup>c</sup> | Total Enterococci (control) | Total *E. coli* (control) | Other *Salmonella* spp. (control) |
| 0 <sup>b</sup> | ND | ND | 5.54 (0.60) | 6.10 (0.85) | <LOD |
| 0.5 | 3.62 (0.35) | 3.67 (1.18) | ND | ND | ND |
| 6 | 4.44 (1.19) | 4.57 (1.38) | ND | ND | ND |
| 24 | 6.44 (0.78) | 6.11 (0.62) | ND | ND | ND |
| 48 | 6.15 (0.05) | 5.53 (0.49) | 6.03 (0.42) | 6.94 (0.42) | <LOD |

Notes.
<sup>a</sup> Abbreviations: SH-2813<sub>nal</sub> – Nalidixic acid resistant *S. Heidelberg*, *E. faecalis*<sub>JH2-2</sub> – Rifampicin resistant *E. faecalis*,
ND – Not determined, LOD – Limit of detection (not detected after 24 h enrichment).
<sup>b</sup> Concentration of *Enterococcus* spp, *E. coli* and *Salmonella* was determined in uninoculated ceca (n =3) for time points 0 and 48 h only.
<sup>c</sup> Each cecum was inoculated with ~ 10<sup>5</sup> CFU of SH-2813<sub>nal</sub> and *E. faecalis*<sub>JH2-2</sub> before incubation under microaerophilic condition.

### Solid plate mating experiments

Bacterial strains used for mating experiments are listed in Table 3. Five colonies from relevant recipient and donor strains (∼ 10^8^ CFU/ml) were selected from overnight cultures grown on sheep blood agar (Remel Inc, San Diego, CA) and resuspended in 900 µl of 1X PBS. Recipient and donor were diluted 10,000 and 100,000-fold respectively, using 1X PBS. Thereafter, 100 µl from the 1:10,000 recipient dilution was spread on CHROMagar and incubated for 6 h. Following incubation, 100 µl from the 1:100,000 donor dilution was spread on to the recipient bacterial lawn and incubated for 18 - 24 h. After incubation, two areas (∼ 0.3 cm) of equal sizes were stamped from the middle and edge of the agar and added to 900 µl of 1X PBS. Vigorous shaking at 1800 rpm was performed with a Fastprep-96 bead beater (MP Biomedicals, Solon, Ohio) for 1 min to suspend the cells from the agar plug. Serial dilutions were then spread on brilliant green sulfur agar (BD Difco, Sparks, MD) or CHROMagar containing relevant antibiotics to distinguish the recipient, donor and “recombinant population” that acquired antibiotic resistance. Agar plates were supplemented with antibiotics at the following concentrations, unless otherwise noted: 8 ppm ampicillin; 1 ppm gentamicin, 16 ppm tetracycline, and 16 ppm streptomycin, all purchased from Sigma (Sigma-Aldrich Corp, St. Louis, MO).

**Table 3.** Resistance phenotype, β-lactamases and plasmids detected in *S.* Heidelberg and *E. coli* donor, recipient and recombinant strains from *in vitro* mating experiments.

| Mating ID <sup>b</sup> | Strain ID | Resistance phenotype <sup>a</sup> | AmpC gene | Plasmids identified | IncK2 contig size (bp) |
| --- | --- | --- | --- | --- | --- |
| **Solid plate** |  |  |  |  |  |
| SH-2813<sub>don</sub> | SH-13A-48h-NalAmp<sup>phe</sup> | Amp, Amc, Fox, Cro, Nal | CMY-2 | IncX1, IncK2 | 88,798 |
| SH-2813<sub>rec</sub> <sup>c</sup> | SH-14A-48h-NalAmp<sup>phe</sup> | Str, Nal | None | IncX1 | NA |
| SH-2813<sub>don</sub> | SH-13A-48h-NalAmp<sup>phe</sup> | Amp, Amc, Fox, Cro, Nal | CMY-2 | IncX1, IncK2 | 88,798 |
| Ec15ceca<sub>rec</sub> | Ec15ceca | Tet | None | IncFIB, Col440i | NA |
| **SH2813_D_Ec15ceca_R** | N/A | Amp, Amc, Fox, Cro, Tet | CMY-2 | IncK2, IncFIB, Col440i | 88,178 |
| SH2813_D_Ec15ceca_R<sub>don</sub> | N/A | Amp, Amc, Fox, Cro, Nal | CMY-2 | IncK2, IncFIB, Col440i | 88,178 |
| SH-2813<sub>rec</sub> | SH-2813-Parental<sup>tyr</sup> | Nal | None | IncX1 | NA |
| **Ec15ceca_D_SH2813_R** | N/A | Amp, Amc, Fox, Cro, Nal | ND | ND | ND |
| Ec6BL<sub>don</sub> | Ec6BL | Amp, Amc, Fox, Cro, Gen, Str, Smx, Tet | CMY-2 | IncK2, IncFIC, IncFIB, IncFII, IncFIA, ColpVC, Col (MG828) | 88,037 |
| SH-2813<sub>rec</sub> | SH-2813-Parental<sup>tyr</sup> | Nal | None | IncX1 | NA |
| **Ec6BL_D_SH2813_R2** | N/A | Gen, Str, Smx, Tet, Nal | None | IncFII, IncX1 | NA |
| **Ec6BL_D_SH2813_R** | N/A | Amp, Amc, Fox, Cro, Nal | CMY-2 | IncK2, IncX1 | 85,244 |
| Ec15BL<sub>don</sub> | Ec15BL | Amp, Amc, Fox, Cro | CMY-2 | IncK2, IncFIC, IncFIB, Col (MG828) | 78,413 |
| SH-2813<sub>rec</sub> | SH-2813-Parental<sup>tyr</sup> | Nal | None | IncX1 | NA |
| **Ec15BL_D_SH2813_R** <sup>d</sup> | N/A | Amp, Amc, Fox, Cro, Nal | CMY-2 | IncK2, IncX1 | 84,268, 84,274, 84,273 |
| **Broth** |  |  |  |  |  |
| SH-2813<sub>don</sub> | SH-13A-48h-NalAmp<sup>phe</sup> | Amp, Amc, Fox, Cro, Nal | CMY-2 | IncX1, IncK2 | 88,798 |
| SH-2813<sub>rec</sub> <sup>c</sup> | SH-2813 | None | None | IncX1 | N/A |

<sup>a</sup> Abbreviations: Amc - Amoxicillin/Clavulanic acid, Amp – Ampicillin, Cro – Ceftriaxone, Fox - Cefoxitin, Gen – Gentamicin, Nal – Nalidixic acid, Smx- Sulfamethoxazole, Str - Streptomycin, Tet – Tetracycline, don – donor, recipient, ND – Not determined, NA – Not applicable.
<sup>b</sup> Boldness denotes recombinant strains with acquired antibiotic resistance.
<sup>c</sup> Recipient did not acquire β-lactamase resistance from donor.
<sup>d</sup> WGS was done on 3 individual colonies that acquired β-lactamase resistance from donor.

### Mating/competition in liquid culture

Mating between SH-13A-48h-NalAmp (SH-2813_don_) and IncK2-free SH-2813 (SH-2813_rec_) was performed in Mueller Hinton Broth (MHB) (BD Difco, Sparks, MD). Donor-recipient mixture was made as previously described (15). Mixtures were diluted 10,000-fold in 50 ml centrifuge tubes containing 20 ml of MHB supplemented with or without 32 ppm amp and incubated under the same microaerophilic conditions as the ceca for 24 h without shaking. To identify SH-2813_rec_ that acquired the IncK2 carrying CMY-2 gene, we screened 56 colonies from each population at 24 h on brilliant green sulfur agar (BGS) supplemented with nal and/or amp. Colonies that grew on BGS supplemented with amp but showing no growth on BGS with nal were designated as “presumptive” recipients of IncK2.

*In vitro* fitness cost associated with acquiring the IncK2 plasmid was tested with 24 h population of SH-2813_don_ and SH-2813_rec_ under no amp selection. The fitness of the population carrying IncK2 plasmids relative to IncK2-free population was determined as described by Miller

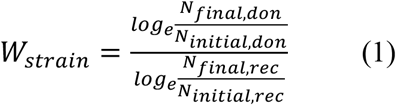

Where *W_strain_* is the fitness of the population carrying IncK2 plasmids (SH-2813_don_), *N_initial,don_* and *N_final,don_* are the numbers of cells (in CFU) in the population carrying the IncK2 plasmids before and after competition, and *N_initial,rec_* and *N_final,rec_* are the numbers of cells of IncK2-free (SH-2813_rec_) population before and after competition. To determine *N_final,don_* and *N_final,rec_* we screened 56 colonies per population from BGS containing no antibiotics onto BGS with nal and amp. From here, the fraction of IncK2 plasmid carrying cells (isolates resistant to both nal and amp) within each population was estimated by scoring the presence of growth on both BGS plates with nal and amp (*N_final,don_*). For IncK2-free population, isolates that were susceptible to amp were used for their estimation (*N_final,rec_*).

### Plasmid copy number determination

We determined the plasmid copy number (PCN) of IncK2 present in each population during liquid culture mating. PCN was determined as described previously (2). Primer sets targeting *gapA* of *S*. Heidelberg and the *inc*RNAI-*rep* of IncK2 were used for PCN determination (Table S1). Real-time qPCR amplification and analysis were performed on a CFX96 Touch Real-Time PCR Detection System (Bio-Rad Inc., Hercules, CA) and carried out as previously described (2). PCN was determined as the copy ratio of plasmid encoded genes to *gapA* (33).

### IncK2 and *bla*_CMY-2_ stability in *S*. Heidelberg

The stability of IncK2 plasmid present in SH-2813_don_ was tested over 50 generations. We serially passed 50 individual colonies of SH-2813_don_ consecutively for 5 days on brilliant green sulfur agar supplemented with and without amp. On the 6^th^ day, hot-boil DNA extraction was performed on 10 µl of each bacterial colony/culture. Afterwards, qPCR was performed on extracted DNA with primers targeting the *inc*RNAI-rep (region upstream of *repA*) of IncK2 and *bla*_CMY-2_ (Fig. 2a; Table S1).

**Fig. 2.**
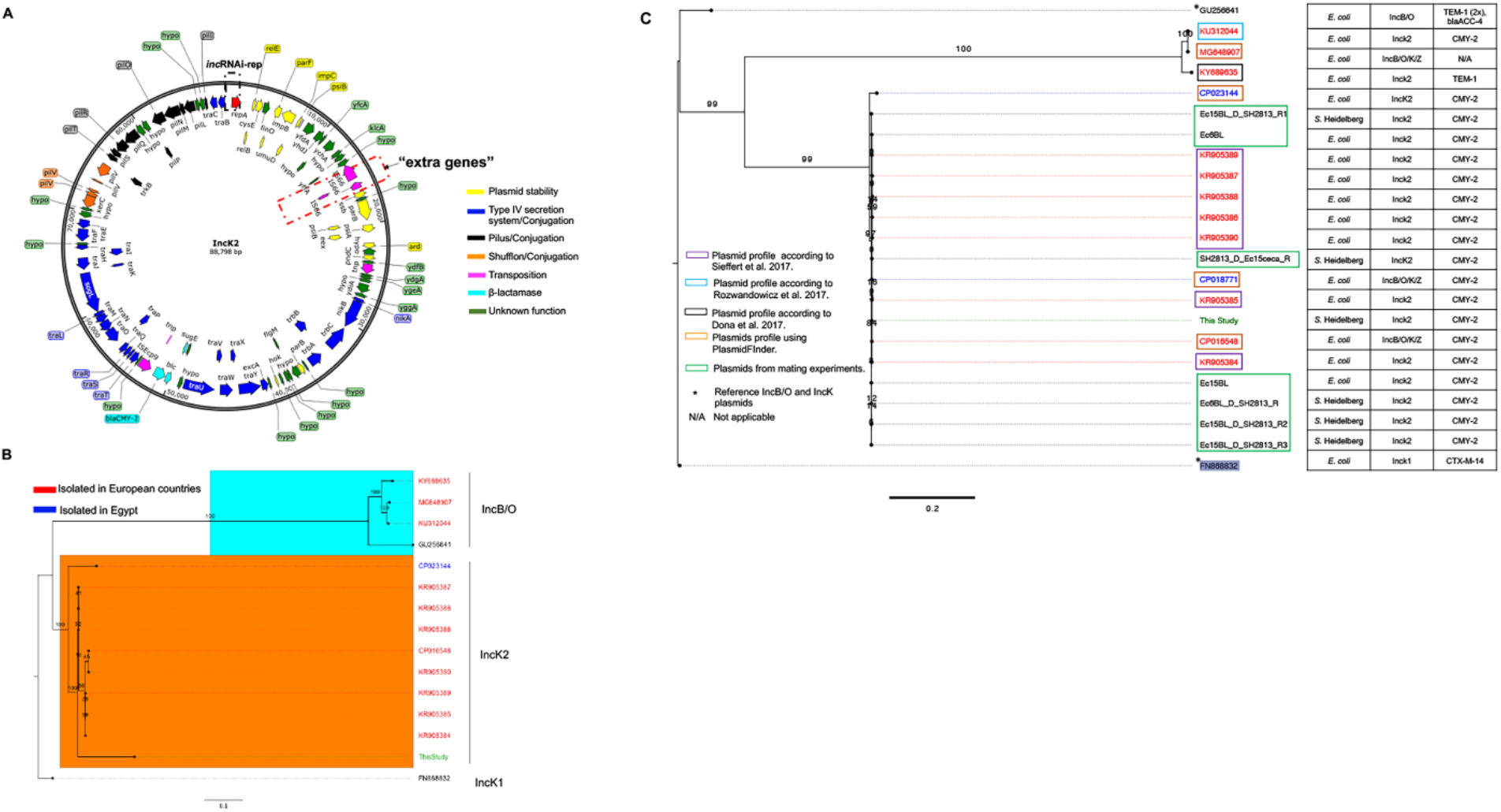
IncK2 acquired by *S*. Heidelberg. (**a**) Annotated map of IncK2 plasmid from this study. Dashed red rectangular box is to highlight “extra genes” discussed in manuscript. (**b**) Maximum likelihood tree of IncB/O/K plasmids based on complete plasmid sequence alignment. (**c**) Maximum likelihood tree of IncB/O/K plasmids based on *inc*RNAi and *rep* gene alignment. The TVM model of nucleotide substitution and GAMMA model of rate heterogeneity was used for sequence evolution prediction. Numbers shown next to the branches represent the percentage of replicate trees where associated taxa cluster together based on ∼100 bootstrap replicates. Trees were rooted with reference IncK1 plasmid (FN868832). Plasmid map was drawn using SnapGene v.4.3.8.1. Tree was viewed using FigTree v1.4.4.

### Statistical analyses

Continuous variables did not meet the assumption of a normal distribution, therefore comparisons between strains or treatment were performed using Wilcoxon signed-rank test. Analyses were performed using R (version 3.4.1).

### Ethics statement

The authors did not physically interact with the chickens before eviscera collection at the processing plant, therefore the authors are exempt from university guidelines and USDA/NIH regulations regarding animal use.

### Data availability

Whole genome sequence and resistome fastq files are available under NCBI Sequence Read Archive (SRA) BioProject: PRJNA509629.

## Results and Discussion

### *In vitro* bacterial survival in broiler ceca

To further explain how mobile elements may contribute to the fitness of *Salmonella* and *Enterococcus*, we monitored the survival of a nal resistant *S*. Heidelberg (SH-2813_nal_) and rif resistant *E. faecalis*_JH2-2_ following microaerophilic incubation in broiler ceca collected from a commercial processing plant. The average cecum weight on collection was 4.4 ± 0.2 g (n =18), with a range of 2.3 – 6.8 g. SH-2813_nal_ and *E. faecalis*_JH2-2_ grew ∼3 and 2 logs after 24 h of incubation in inoculated ceca. No *Salmonella* was detected in uninoculated ceca, however total *E. coli* increased by ∼1 log after 48 h of incubation (*p*.value < 0.086; Wilcoxon signed-rank test). No significant change was observed in Enterococci populations in uninoculated ceca (Table 1), but low background rif resistant populations were detected on Enterococcosel agar supplemented with rif.

Although SH-2813_nal_ and *E. faecalis*_JH2-2_ were able to grow *in vitro* in ceca, this does not correspond to colonization – attachment of bacteria to ceca epithelial cells. For instance, Mchan et al. (34) demonstrated *in vitro* that up to 5 logs of *S*. Typhimurium could successfully attach to the ceca of 1 and 2 week old broiler chicks. In our study, the final *S*. Heidelberg concentration measured in ceca was log 6.15 ±0.05 CFU/g and likely includes attached and unattached bacterial cells.

### Acquired antibiotic resistance after 48 h incubation in ceca

To determine if *E. faecalis*_JH2-2_ acquired AMR following 48 h incubation in the cecum of broilers, we performed AST on selected isolates (n = 9) recovered during incubation. No *E. faecalis*_JH2-2_ isolate acquired resistance to any antibiotic present on the NARMS Gram-Positive panel (data not shown), therefore *E. faecalis*_JH2-2_ will not be discussed further in this study. To determine if SH-2813_nal_ acquired resistance to β-lactam antibiotics, we used XLT-4 agar supplemented with nal and amp to select for presumptive isolates. In addition, AST was done on presumptive β-lactam producing isolates (n = 4) and a selected number of SH-2813_nal_ isolates recovered from XLT-4 agar with nal only (n = 9). We chose β-lactam due to the reported increase in cephalosporin resistance in *S*. Heidelberg recovered from chicken and chicken products (5, 35). Five of the thirteen isolates including the 4 presumptive β-lactam isolates acquired increased resistance to multiple antibiotics after 48 h of incubation of ceca. Two isolates displayed complete or intermediate resistance to cefoxitin (β-lactams), chloramphenicol (phenicols) and ciprofloxacin (fluoroquinolones) and 2 other isolates were resistant to tetracyclines (streptomycin and/or tetracycline), sulfizoxazole (sulfonamides) and cefoxitin (Table 2). One isolate (SH-13A-48h-NalAmp) showed decreased susceptibility or resistance to multiple β-lactams including cefoxitin, ceftriaxone, cefazolin, cefpodoxime, ceftazidime, cefuroxime, piperacillin, ticarcillin/clavulanic acid, ampicillin, ampicillin/sulbactam and amoxicillin–clavulanic acid. AST was performed twice on these 5 isolates to ascertain that observed changes in MIC were not reversed after cultivation.

**Table 2.**
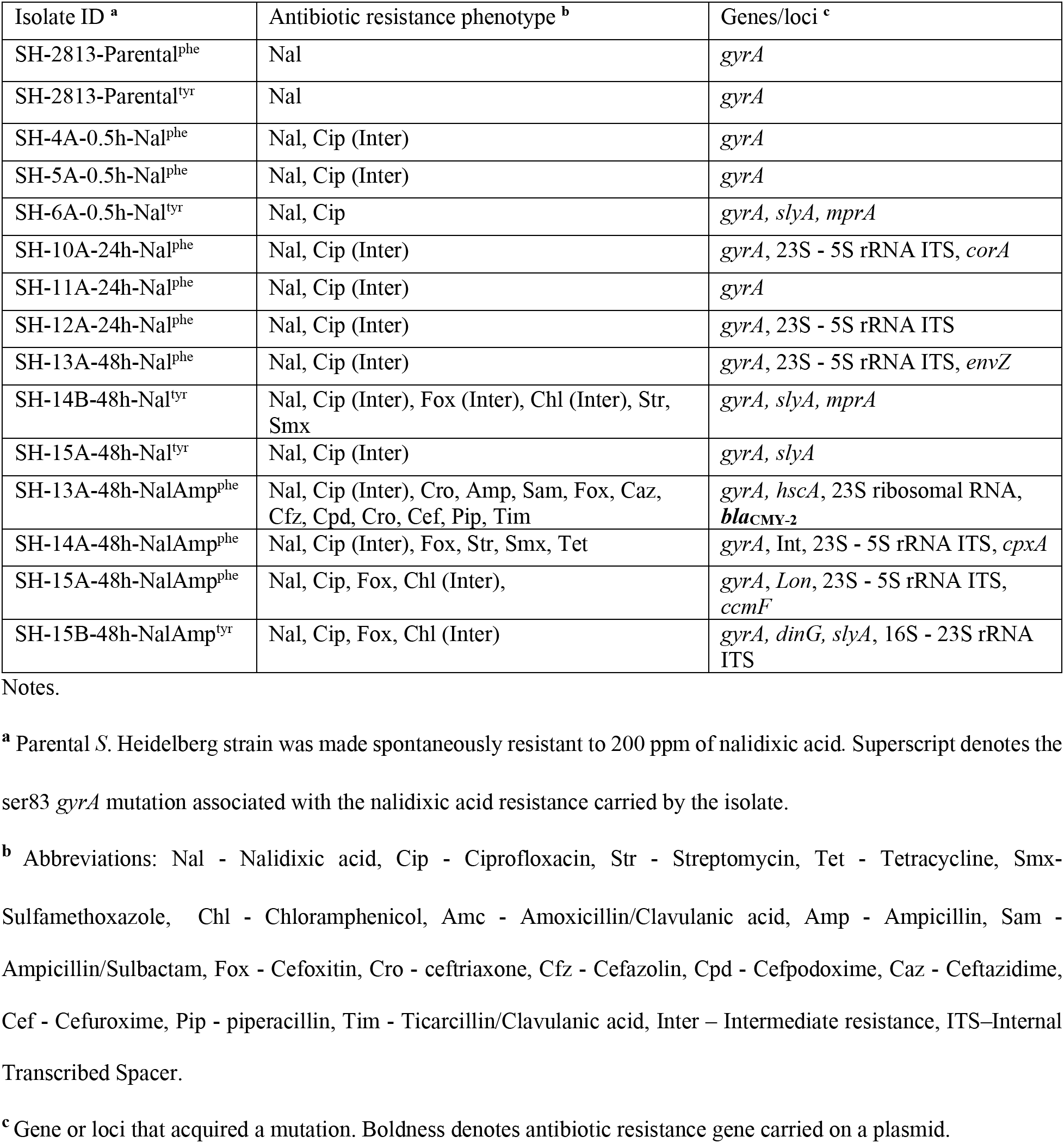
Acquired antibiotic resistance profile of *S*. Heidelberg isolates recovered after ceca incubation.

To identify the genomic changes underlying the acquisition of antibiotic resistance in these 5 isolates, WGS data from all isolates including sensitive and resistant isolates were compared using SNP/indel and resistance gene identification. Each of the resistant isolate carried at least one unique mutation from other sensitive/resistant isolates and the reference genome, except isolate SH-14B-48h-Nal (Table 2 and Table S2). These mutations were in phage integrase (*int*), sensor histidine kinase (*cpxA*), ATP-dependent serine endopeptidase La (*lon*), cytochrome c-type biogenesis protein (*ccmF*), ATP-dependent DNA helicase (*dinG*), iron sulfur protein assembly (*hscA*) and the 16S-23S rRNA Internal transcribed spacer (ITS) regions of the genome (Table S2). Among these loci, only the *cpxA* gene has been reported to confer resistance to cephalosporins via transcriptional regulation of multiple efflux pumps such as OmpC and OmpF (36, 37). This mutation was identified in only one isolate. The observed phenotypic resistance could not be matched with a known mutation for the other four isolates with acquired resistance. For quinolones, *gyrA* mutations have been shown to confer cross-resistance to nalidixic acid and ciprofloxacin (38). Here, the parental strains (SH-2813_nal_) had either a Ser83tyr or Ser83phe substitution in *gyrA* which could have accounted for the increase in MIC for ciprofloxacin in all isolates tested.

A β-lactamase gene (*bla*_CMY-2_) was found in 1 isolate (SH-13A-48h-NalAmp) that was likely responsible for the observed β-lactam resistance phenotype in this isolate. The *bla*_CMY-2_ was located on a contig totaling 88 kb and predicted as IncB/O/K/Z plasmid using PlasmidFinder. The newly acquired plasmid had a read coverage two times greater than the chromosome and harbored stability, conjugative transfer and shufflon recombinase genes (Fig. 2a), suggesting a plasmid with ∼2 copies per chromosome. Recently, Roschanski et al. showed that an *E. coli* isolate carrying an IncB/O/K/Z-like plasmid encoding a *bla*_CMY-2_ gene conferred resistance to cephalosporins and carbapenems and hypothesized that an overexpression of CMY-2 and high plasmid copies (8 per cell) was responsible for decreased susceptibility to carbapenems (39). Here, this isolate exhibited increased resistance to multiple cephalosporin and penicillin drugs but was susceptible to the carbapenems tested (Table 2), indicative of extended-spectrum β-lactamase (ESBL)-like producing *S*. Heidelberg strain.

For antibiotic resistant isolates without a known ARG it is possible that over expression of multidrug efflux pumps such as *acrAB-tolC*, *marA sdiA*, *mdsA*, *mdsC* etc. contributed to the higher MIC for cefoxitin, chloramphenicol and tetracyclines. These genes were present in parental SH-2813_nal_ (Table S3) and a up/down regulation of these efflux pumps have been shown to confer resistance to various antibiotic classes. For instance, Suzuki et al. demonstrated AR acquisition in *E. coli* could be quantitatively predicted from the expression changes of a few genes such as *OmpF* and *acrB* (36). No resistance gene or mutation was identified to be associated with sulfizoxazole resistance, which agrees with other studies that have reported the lack of sulfonamide resistance genetic elements in some *Salmonella enterica* isolates (40, 41).

The harsh chicken gut environment may induce proteins that could confer cross protection against antibiotics. In a recent publication, the authors exposed chicken breast inoculated with 15 outbreak-linked *Salmonella enterica* strains to simulated gut conditions of the mouth, stomach and intestines (42). Acquisition of resistance to ciprofloxacin, ampicillin, ceftriaxone and sulfamethoxazole-trimethoprim was reported for isolates of serovar Heidelberg, Newport, Albany and Corvallis after 1 h of exposure.

### Comparative genomics of IncB/O/K/Z-like plasmid acquired from ceca

To compare the IncB/O/K/Z-like plasmid found in SH-13A-48h-NalAmp with sequenced IncB/O/K/Z plasmids, we sequenced plasmid DNA extracted from this isolate. An assembly of the sequenced genome with SPAdes resulted in 2 putative plasmid contigs of 88,798 and 37,845 bp sizes with a coverage > 200X. The completeness and circularity of these plasmid contigs was inferred as described previously (2). The 37.8 kb plasmid is an IncX1 plasmid carried by the parental SH-2813_nal_ and was present in all *S.* Heidelberg sequenced for this study. Querying the NCBI nr sequence database with the 88 kb contig yielded plasmids belonging to the IncK2 compatibility group (43) as the closest homologs (data not shown). These plasmids carried the *bla*_CMY-2_ gene and ranged in size from 79 – 86 kb. These IncK2 plasmids were found in *E. coli* strains isolated from clinical and poultry samples collected in European countries (43, 44). A phylogenetic tree reconstructed with selected complete IncK2 and IncB/O DNA sequences showed that the plasmid from this study shared up to 62 % pairwise identity with IncK2 plasmids reported by Sieffert et al. al. (44) and < 25 % identity with plasmids of the IncB/O compatibility group. Consequently, all IncK2 plasmids were represented by a separate clade on the reconstructed ML tree (Fig. 2b).

The IncB/O, IncK, and IncZ plasmids belong to the I-complex of plasmids and share high homology in their incompatibility RNAI sequences (non-coding antisense RNA that is upstream of *ori*) (43, 45, 46). Therefore, the use of the *inc*RNAI region as targets in the plasmid-based replicon typing (PBRT) classification scheme of the IncB/O/K plasmids poses challenges with their typing (43). Here, a phylogenetic tree reconstructed using the *inc*RNAI-rep sequences from plasmids sharing significant *inc*RNAI region homology (e-value < 0.0001) with the plasmid from this study showed that this plasmid is more closely related to the IncK2 plasmids (Fig. 2c) than to the IncB/O group. This supports the result from the whole plasmid DNA comparison. Based on this comparative analysis, this IncB/O/K/Z-like plasmid can be assigned to the IncK2 subgroup of IncI-complex plasmids.

The major difference between the IncK2 plasmids was in the coding DNA sequence (CDS) regions for shufflon proteins (Fig. S1) which has been previously reported (44). However, the IncK2 from this study carried three extra IS66-family transposases totaling ∼ 2.5 kb inserted between the single-stranded DNA-binding protein (*ssb*) and an uncharacterized protein (*yffA*) (Fig. 2a and Fig. S1).

### *In vitro* stability, fitness and plasmid copy number of IncK2 in host

We serially passed 50 colonies from the SH-2813_don_ on brilliant green sulfur agar with or without amp for 50 generations and used qPCR primers targeting the *inc*RNAI-rep region of IncK2 to confirm their presence. The IncK2 plasmid exhibited 100 % stability under amp selection and 86% stability without selection. In contrast, the CMY-2 gene was only stable in 25/50 colonies (50 %) under amp selection and 23/50 (46 %) without selection. Rozwandowicz et al. also reported 100 % stability for IncK2 plasmids carried by *E. coli* isolates from Netherlands after 30 generations in LB agar supplemented with and without antibiotics (43). The temporal stability of *bla*_CMY-2_ gene has been previously reported for strains of *Salmonella* serovar *enterica* (47).

Next, we performed mating experiments in broth culture to simultaneously assess the transfer, fitness cost and PCN control of IncK2 in SH-2813_don_. This experiment was done with SH-2813_don_ and IncK2-free SH-2813_rec_ in the presence or absence of ampicillin selection. Transfer of IncK2 to SH-2813_rec_ was not observed in broth experiments. Nevertheless, IncK2-carrying SH-2813_don_ population had significantly lower fitness than IncK2-free SH-2813_rec_ population during mating experiment under no selection (*p*.value = 0.0636; Wilcoxon signed-rank test, Fig. 3b). The PCN of IncK2 was moderate to high during the mating experiment ranging from 6 to 15 PCN/cell under selection and 21 to 35 PCN/cell without selection (Fig. 3a). This suggests that during ampicillin exposure IncK2 is present at low copies per cell but in the absence of ampicillin IncK2 is present at high copies per cell.

**Fig. 3.**
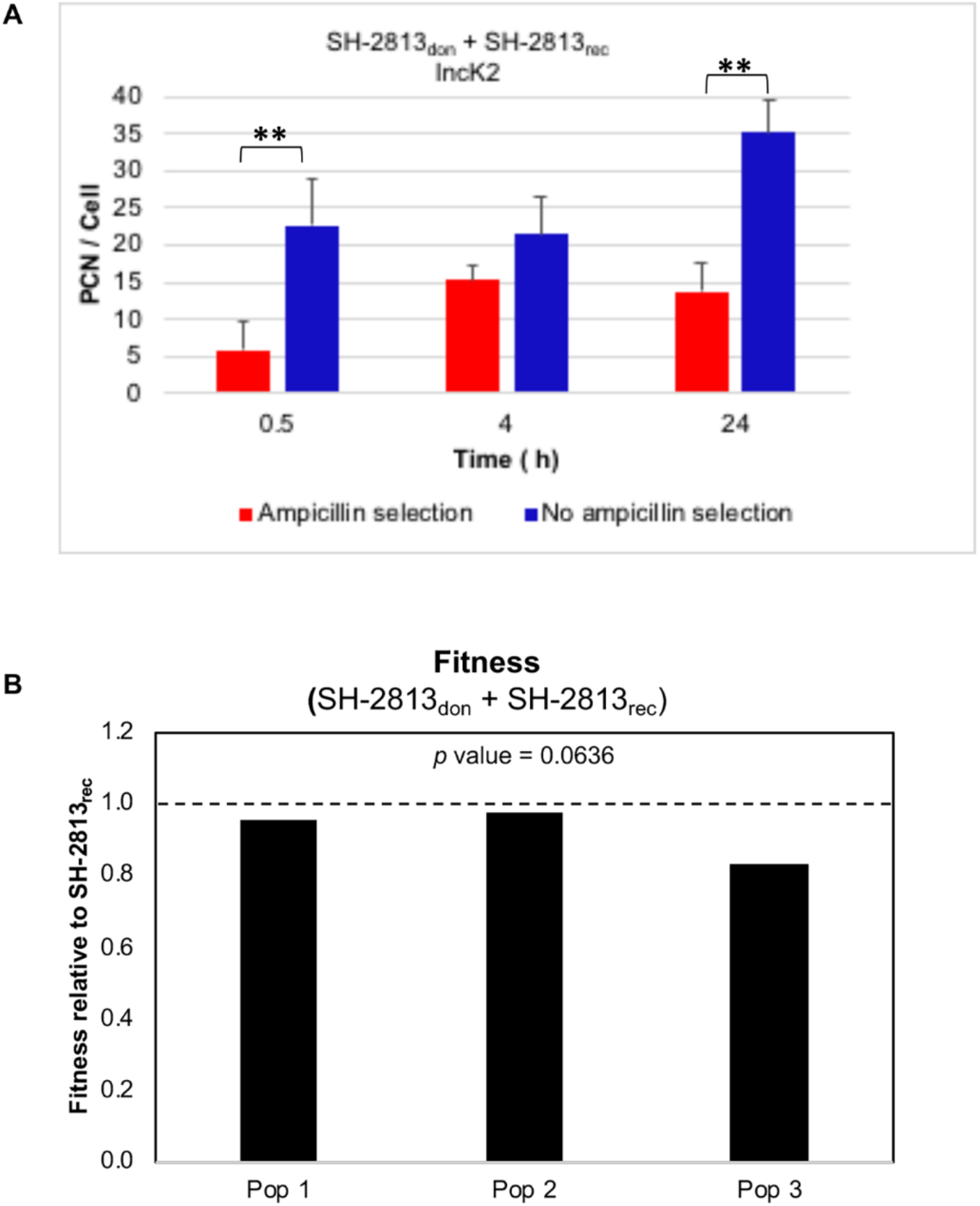
*In vitro* plasmid copy number (PCN) and fitness of IncK2 in *S*. Heidelberg host. IncK2 PCN (**a**) was determined during mating experiment between IncK2-carrying populations (SH-2813_don_) and IncK2-free SH-2813_rec_ populations with and without ampicillin selection (**b**) Fitness of IncK2-carrying populations relative to IncK2-free populations under no ampicillin selection. Each bar represents the fitness of one population that was established from one single bacterial colony. The horizontal dashed line represents the fitness of SH-2813_rec_.

### Relative abundance of β-lactamase genes and *Proteobacteria* in ceca

We performed targeted enrichment of ARG’s using hybridization capture of shotgun metagenomic library. Our goal was to sequence ARG’s at a higher depth of coverage and determine the relative abundance of *bla*_CMY-2_. This enrichment approach has been demonstrated to increase ARG detection several fold compared to shotgun metagenomics (48). Here, we validated this method using WGS and phenotypic data on MDR pathogens (see supplemental methods). The average coverage of de novo assembled contig carrying an ARG was 803 X with a range of 11 – 32735 X. This method showed a high level of concordance with our WGS dataset and all false positives were removed with a coverage threshold of 10X (Table S4, Fig. S2).

The most abundant ARG’s in ceca upon collection confer resistance to aminoglycosides, tetracyclines macrolides, lincosamides or streptogramins class of antibiotics (Fig. 4). Following the inoculation of *S*. Heidelberg into ceca, ARG’s decreased in abundance and a few ARG’s that were undetected at time zero were detected at a later time point (Fig. 4). For instance, *aadA1* and *tetA* genes were not detected until after 6 h of incubation. Both genes are commonly found on class 1 integrons carried by *Enterobacteria* (49, 50) suggesting that facultative anaerobes including *E. coli* may be blooming *in vitro*. Four β-lactamase genes were detected at low abundance but *bla*_CMY-2_ was not detected at any time point (Fig. 4). Two genes (*cfxA* and *cepA-49*) found mainly in strict anaerobes including the *Bacteriodetes* (51, 52) were the β-lactam genes detected at time zero, which is indicative of the microaerophilic condition of the ceca upon collection. The *bla*_TEM-1B_ gene that is commonly found in β-lactamase resistant *E. coli* (53) was the only other β-lactam gene detected (Fig. 4a). However, *bla*_TEM-1B_ was not detected until time point 6 h. Collectively, the resistome data suggest that the condition in the ceca may be getting less anaerobic *in vitro*, thus favoring an increase in ARG’s harbored by facultative anaerobes. This is further corroborated by our culture data that revealed a log increase in *E. coli* population after 48 h of ceca incubation (Table 1).

**Fig. 4.**
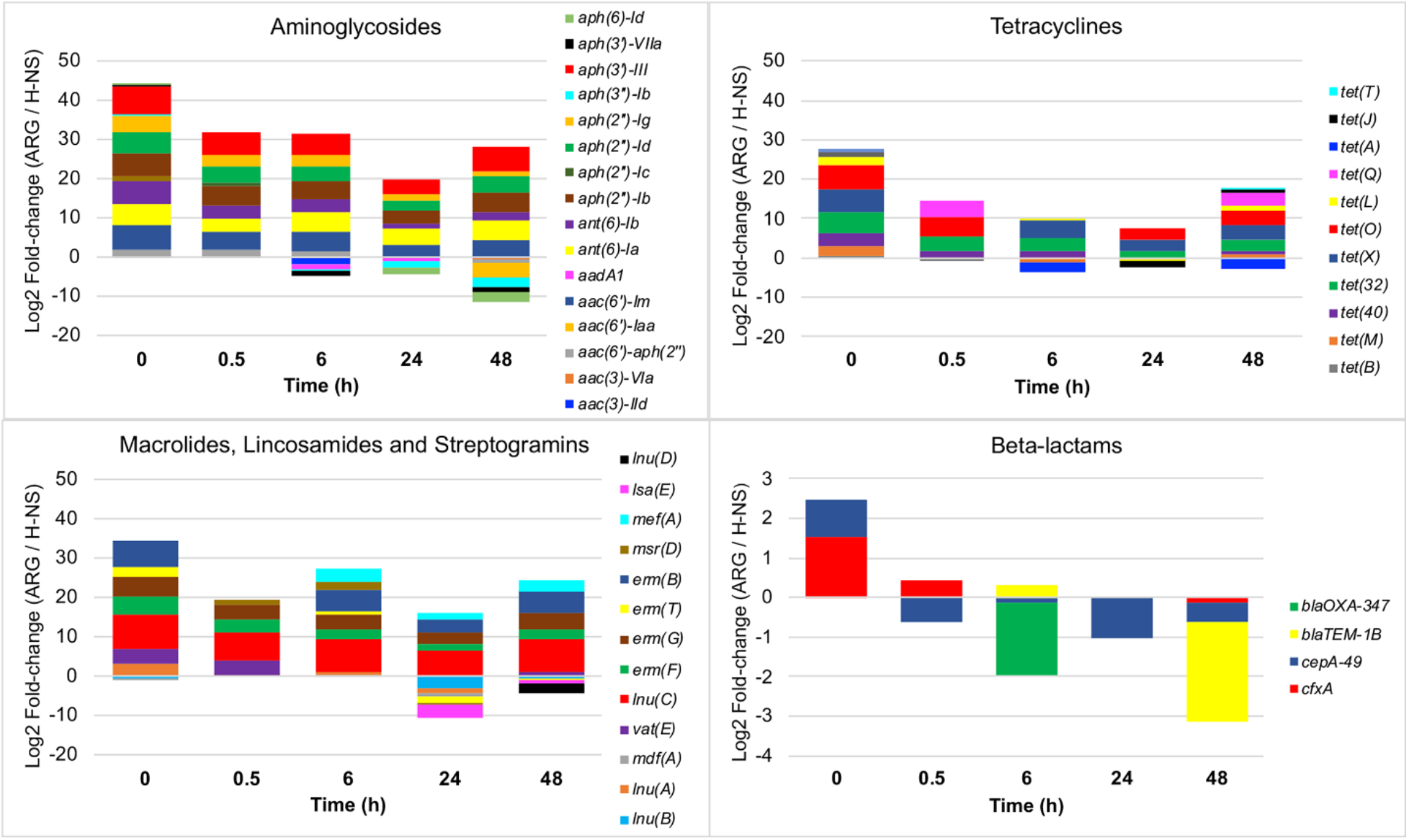
Relative abundance of ARG’s determined using resistome enrichment. Targeted enrichment of ARG’s using hybridization capture of shotgun metagenomic library was done on each cecum. At each time point, three ceca samples were used for resistome determination and ARG abundance was calculated by dividing the ARG contig coverage by the coverage of H-NS. Positive value indicates ARG abundance is higher than H-NS while a negative value indicates ARG abundance is lower than H-NS.

For the broiler ceca used for this study, the *bla*_CMY-2_ gene represented a negligible fraction of their resistome. The *bla*_TEM-1B_ contig had up to a 200X coverage indicating that the enrichment method should also be sufficient for the detection of *bla*_CMY-2_. One explanation for the non-detection could be the instability of *bla*_CMY-2_ as we discussed previously. Furthermore, preserving the cecal contents in RNAlater prior to resistome enrichment may have introduced some artifacts. Although biases associated with microbiome and transcriptomic analyses of samples stored in preservatives have been reported (54), we do not know their effects on plasmids or ARG stability.

### Retrospective identification of potential IncK2 donors

The “extra” genes observed in the IncK2 plasmid from this study were part of a ∼ 9 kb region identified as P1 bacteriophage SJ46 (NC_031129) by PHAST (Fig. S3). This region encoded CDS annotated as entry exlusion protein (*eex*) alleviation of restriction of DNA (*ard*), plasmid stability genes and IS66 family of transposons. This suggests that IncK2 plasmid may be a chimera of P1-like phages with an IncK2 plasmid backbone. In addition, the *repL* representative gene of the P1 phages was detected in two of the IncK2 reference plasmids (Fig. S1). It is unlikely that this IncK2 plasmid was transferred from another *Salmonella enterica* strain because no *Salmonella* with IncK2 to our knowledge has been characterized or reported. Also, the ceca samples used in this study were negative for *Salmonella* even after 24 h enrichment in buffered peptone water. Additionally, NCBI BLAST search against the nr database found the closest matches were to plasmids in *E. coli* strains suggesting *E. coli* was a potential donor of this plasmid. Based on this premise, we retrospectively screened 16 *E. coli* isolates recovered from 6 ceca samples for IncK2 plasmids.

Only 1 of the 16 isolates (hereafter referred to as Ec15ceca) tested positive for IncK2 via qPCR. However, after WGS this isolate harbored no IncK2 plasmid but carried IncFIB and Col440i plasmids, and a *tetA* gene that conferred tetracycline resistance (Table 3). This made us question if qPCR primers targeting the *inc*RNAI-rep region of IncK2 plasmids (43, 44) were amplifying homologous regions in the genome of Ec15ceca. To answer this question, we performed a BLAST (tblastx) search against the de novo assembled genome of Ec15ceca with the *inc*RNAI-rep (∼1,476 bp) sequence from plasmids denoted IncK2 in Fig. 2b. The only significant match (*e*.value < 0.02) was a ∼ 11.7 kb contig with ∼ 39 % pairwise identity. Protein annotation of this contig predicted 11 of 13 CDS to encode phage tail assembly proteins (Fig. S4). This suggests that the putative phage with this contig may have packaged plasmid DNA during exit from a bacterial cell carrying IncK2 (55). Additionally, this tail module may encode variable regions that determine the specificity of phage attachment to different bacterial hosts (56).

Notwithstanding the limitations of phage genome assembly using Illumina short read technology, we used phage search tools to identify phage DNA present in the genome of Ec15ceca and two *E. coli* isolates carrying IncK2 collected from broiler litter (Table 3). PHAST and PHASTER identified varying number of putative phage contigs, this included P1-like phages that were identified multiple times in queried contigs (Table S5). The largest contig identified carrying the required modules for P1 virion assembly (57) was ∼ 113 kb (Fig. S5). This phage encoded 2 of the 3 “extra” IS66-family transposases present in the IncK2 plasmid from this study. Lastly, a positive result with qPCR primers targeting the *repL* gene (58) corroborated the presence of a P1-like phages in these *E. coli* strains.

Together, the results suggest that a P1-like phage present in *E. coli* and an IncK2 plasmid may have been involved in a recombination event during ceca incubation. P1 phages were first identified in the 1950’s and have been extensively reviewed and studied (57). They are bacteriophages of *Enterobacteriaceae* that lysogenize their host as circular plasmids but are active virions when in lytic phase. P1 phages can package with them large DNA material with them upon exit from their host, making them a workhorse for genetic exchanges via generalized transduction (57). Also, P1 phages carry antirestriction and restriction modification genes that help them evade bacterial host killing by restriction (Fig. S5). Consequently, P1-like phages may facilitate the transfer of plasmids and ARG to new bacterial hosts. For example, P1-like bacteriophages carrying *bla*_CTX-M-27_, *bla*_CTX-M-15_, *bla*_SHV-2_ and *mcr-1* have been reported in *Salmonella* and *E. coli* strains (58–61).

### IncK2 plasmid is transferable between *E. coli* and *S*. Heidelberg

We made attempts to recreate the recombination event between Ec15ceca and SH-2813_don_ *in vitro* using a solid plate mating. Our goal was to determine if transfer was possible, rather than determining the rate of transfer. First, we used Ec15ceca as the recipient of IncK2 and SH-2813_don_ as the donor. After 24 h of mating, we observed multiple *E. coli* colonies on CHROMagar plates supplemented with ampicillin. Whole genome sequencing and AST on a selected colony confirmed the successful transfer of IncK2 carrying *bla*_CMY-2_ to Ec15ceca (Table 3). Next, we performed the same experiment with the Ec15ceca strain that acquired IncK2 (Ec15ceca_don_) serving as the donor to SH-2813_rec_. Here, we observed two SH-2813_rec_ colonies that acquired β-lactam resistance. In addition, we performed mating experiment between SH-2813_don_ and SH-2813_rec_ but observed no transfer of β-lactam resistance.

Finally, we selected two *E. coli* strains isolated from the broiler litter (BL) of a grow-out house carrying *bla*_CMY-2_ on an IncK2 plasmid. Strain Ec15BL carries only *bla*_CMY-2_ gene, whereas Ec6BL carried additional ARG’s on other type of plasmids (Table 3). The IncK2 in Ec6BL also carries three “extra” IS66-family transposases (data not shown). We performed the same solid mating experiment with these two strains acting as the donor of IncK2 to SH-2813_rec_. For both strains, we observed transfer of the IncK2 to SH-2813_rec_ after 24 h of mating (Table 3). A plasmid designated IncFII carrying multidrug resistance and virulent genes was also transferred from Ec6BL to SH-2813_rec_ (Fig. S6).

Although de novo assembly of plasmids from recombinants resulted in incomplete contigs, protein annotation revealed that the core genome of IncK2 did not change from donor to recipient (data not shown). Also, the *bla*_CMY-2_ backbone (*ISEcp9*-*bla*_CMY-2_-*blc*-*sugE*) was conserved in all recombinants. The IncFII contig (∼ 119 kb) carried ARG’s that conferred resistance to aminoglycosides, tetracyclines and sulphonamides (Fig. S6) and was calculated to be present at ∼4 copies/cell. In addition, this contig carried genes for metal resistance (*mer* operon), colicin production and multiple toxin-antitoxin systems. Interestingly, a ∼38.8 kb segment of the IncFII contig was identified as phage SJ46 and it encoded a class 1 integron (*intI1*), transposases and the aforementioned ARG’s. Also, this IncFII plasmid shared significant homology (*e*.value < 0.0001) with the *inc*RNAI-rep region of IncK2 plasmids.

Determining the mode of recombination responsible for the transfer of these plasmids between *E. coli* and *S*. Heidelberg is beyond the scope of the study, but our analysis suggests that phage-mediated transfer cannot be ignored. The detection of P1-like phage DNA in IncK2 and IncFII genomes and in the core genome of *E. coli* strains suggests that the interactions between phage-plasmid and host are complex. Our hypothesis is that P1-like phages including SJ46 transduce plasmids between permissive hosts (generalized transduction). This does not preclude plasmid transfer that has been shown to occur through the uptake of free DNA (transformation) made possible after viral lysis of host cell (62). However, *Salmonella* has not been demonstrated to be naturally competent.

For conjugation to occur, cell-to-cell contact is required i.e. donor bacterium and recipient bacterium will need to meet. Although this is achievable *in vitro*, it is a bigger challenge in the ceca. One limiting factor to finding a mate carrying IncK2 is the low relative abundance of *Proteobacteria* including *E. coli* and *Salmonella* residing in the chicken gut (7, 63). In contrast, lysis of an *E. coli* strain carrying a P1-phage can produce up 200 infective phage particles per cell (57) that are capable of generalized transduction (64). The rate of transduction will depend non-linearly on the turnover and replication rates of *E. coli* and *Salmonella* in the ceca (64).

The inability to transfer IncK2 between Heidelberg strains *in vitro* was unexpected. Zhang and LeJeune (65) demonstrated that phage-mediated transfer of *bla*_CMY-2_, *tetA* and *tetB* genes from *S*. Heidelberg to *S*. Typhimurium was possible. One explanation is that the conditions used *in vitro* for mating experiments in our study are not reflective of that in the ceca. Otherwise, transfer may require phage or plasmid genes that are only available when *E. coli* is present.

Increasing resistance to cephalosporins has been attributed to the production of plasmid-mediated extended-spectrum or AmpC β-lactamases. The majority of these β-lactamase genes (*bla*_CMY-2_ and *bla*_CTX-M_) have been found on the IncI complex plasmids (35, 66–68). In this study, we show that *S*. Heidelberg could acquire *bla*_CMY-2_ carried on an IncK2 plasmid from *E. coli*. Our analysis of the IncK2 plasmids suggests that PBRT is not sufficient for their detection or discrimination. Consequently, PCR primers targeting the *inc*/*ori* sites of IncK2 are likely to detect homologous regions present in P1-like phages or IncFII plasmids. These results highlight the complex nature of interactions between plasmid-phage and ARG transfer between commensals and pathogens in the gut and supports the need for a more targeted approach.

## Conclusion

Although it is important that *S*. Heidelberg acquired IncK2 carrying *bla*_CMY-2_ *in vitro*, however, this result must be appraised within the context of the experimental design employed. For this same dynamic to occur *in vivo*, a permissive Heidelberg strain will have to evade the physical and biological barriers of the gut to get to the cecum where a bacterial strain carrying *bla*_CMY-2_ must be a co-resident. Furthermore, the inability to transfer IncK2 between Heidelberg strains may result in a bottleneck of IncK2 plasmid populations *in vivo*. The latter hypothesis still needs to be tested. Instead, this study should be viewed as new information on HGT and acquisition of multidrug resistance in *S*. Heidelberg.

## Acknowledgements

We are grateful to Denice Cudnik, Dr. Mark Berrang, Steven Knapp, Carolina Hall, Mary Katherine Crews and Anna-Marie Bosch for their logistical and technical assistance. Any opinions expressed in this paper are those of the authors and do not necessarily reflect the official positions and policies of the USDA and any mention of products or trade names does not constitute recommendation for use. The authors declare no competing commercial interests in relation to the submitted work.

---

**Text S1.** Supplemental materials and methods.

**Table S1.** Primers used for this study.

**Table S2.** Mutations acquired by *S*. Heidelberg isolates following incubation in broiler ceca

**Table S3.** Antibiotic resistance genes and efflux pumps encoded on the chromosome of *S*. Heidelberg.

**Table S4.** Concordance between WGS and resistome enrichment for ARG detection in MDR pathogens.

**Table S5.** Phages identified by PHAST in *S.* Heidelberg and *E. coli* donor and recipient strains used in *in vitro* mating experiments.

**Fig. S1** ProgressiveMauve alignment of IncK2 complete plasmid DNA sequences from this study and Seiffert et al., DNA regions that differ between the IncK2 from this study and Seiffert et al. are highlighted with dashed horizontal black rectangular boxes.

**Fig. S2** Correlation between WGS and resistome enrichment-based relative abundance determination. ARG abundance was calculated by dividing the ARG contig coverage by the coverage of H-NS. ARG’s (n =31) present in MDR *Salmonella* serovars (n =5) was used for Kendall tau’s correlation test.

**Fig. S3** Annotated map of predicted P1-like phage genome present in the IncK2 plasmid from this study. Dashed diagonal black lines denotes phage SJ46 region identified by PHAST. Map was drawn using SnapGene v.4.3.8.1.

**Fig. S4** Annotated contig of strain Ec15ceca sharing homology with IncK2 *inc*RNAi-rep region. The *inc*RNAi-rep sequence of IncK2 plasmids was queried (tblastx) against the de novo assembled genome of Ec15ceca. Map was drawn using SnapGene v.4.3.8.1.

**Fig. S5** P1-like phage identified in *E. coli* genome by PHAST. Dashed red rectangular box is to highlight “extra genes” discussed in manuscript.

**Fig. S6** Annotated map of putative IncFII contig (119.056 bp) transferred *in vitro* from *E. coli* to *S*. Heidelberg. Dashed rectangular red box denotes “predicted” P1-like phage region identified by PHAST. Map was drawn using SnapGene v.4.3.8.1.

